# Age dependent collective trafficking of tissue resident T cells in zebrafish

**DOI:** 10.64898/2026.07.14.738523

**Authors:** Udochi F. Azubuike, Patricia B Gordon, Khanh Loan Ly, Woong Young So, Lanie Le, Kevin Bishop, Raman Sood, Michael Kruhlack, Michael M. Gottesman, Kandice Tanner

## Abstract

How tissue-resident T cells are organized to provide immune surveillance in healthy peripheral tissues and how this organization changes with age remains largely unknown. Here we use intravital imaging in zebrafish to identify a previously undescribed, appendage-specific mode of adaptive immune organization: fin-resident T cells undergo coordinated collective “streaming” migration within connective tissue compartments. T cells colonize the developing fin prior to lymphatic or blood vessel formation, indicating that initial residency can be established independently of classical immune conduits. However, collective streaming is not immediate; it emerges during juvenile maturation, coincident with maturation of fin architecture, and is restricted to T cells rather than other leukocyte populations. After fin amputation, young adults restore streaming within regenerated tissue once spatial compartments are re-established, whereas middle-aged fish following thymic involution repopulate the fin yet fail to promptly recover coordinated streaming and show broader tissue dispersion and altered motility in comparable times. Transcriptomic profiling across ages, regenerative states, and immune-altered microenvironments reveals coupled regulation of stromal remodeling programs and antigen presentation pathways, including differential expression of MHC class I and II components with strong microenvironmental dependence. These data define an age-regulated, tissue-instructed program that organizes resident T cell dynamics and immune potential in situ, providing a framework for understanding how tissue specific stromal environments constrain immune surveillance.

**Significance Statement:** T cells patrol most organs, but we know surprisingly little about how they are arranged inside healthy tissues over an animal’s lifetime. By imaging zebrafish in vivo, we discovered that fin-resident T cells do not move independently: they organize into coordinated “streams” that migrate collectively through fin connective tissue. This collective behavior appears during juvenile maturation, re-forms after regeneration in young adults, and is delayed or disrupted in older fish as age-related thymus decline reduces new T cell production. In parallel, aging and the tissue microenvironment reshape the expression of genes involved in antigen presentation. These results reveal a tissue-specific, age-regulated architecture for adaptive immune surveillance that links aging to impaired immune organization during regeneration.

## Introduction

T cells are central components of the adaptive immune system and are distributed throughout nearly every organ and tissue in mammals and many vertebrates (1, 2). This broad tissue residency enables continuous immune surveillance and rapid responses to infection, injury, and malignant transformation (3). Importantly, immune function within each tissue is not static but varies with developmental stage, age, and environmental history, including prior infections and toxin exposure (1, 3–5). These factors contribute to heterogeneity in T cell responses, influencing cytotoxic potential, tolerance, and susceptibility to disease, and are increasingly recognized as determinants of immunotherapy efficacy in aging populations (4–8).

Despite extensive knowledge of T cell development and activation, far less is understood about how T cells are organized and maintained within peripheral, non-lymphoid tissues over the lifespan (8). In mammals, hematopoietic stem cells give rise to lymphoid progenitors that migrate to the thymus, where T cells undergo maturation and selection before dispersing to secondary lymphoid organs and peripheral tissues (9, 10). Barrier tissues such as skin and gut represent major sites of T cell residency, where local microenvironments shape immune behavior (9, 11). However, new T cell output declines dramatically following the evolutionarily conserved process of thymic involution. How peripheral, tissue-resident T cells are spatially organized, functionally maintained, and remodeled after thymic involution remains a fundamental unresolved question in immunology.

Zebrafish provide a powerful vertebrate model to address these questions. Core features of adaptive immunity, including T cell developmental pathways, migratory cues, and functional subsets, are conserved between zebrafish and mammals (12–17). Thymic involution also occurs in zebrafish at ages comparable to those in humans (18, 19). In addition, zebrafish offer unique experimental advantages: optical transparency enables direct, long-term visualization of immune cell behavior in vivo, and the remarkable regenerative capacity of the fin provides a tractable system to study immune dynamics during tissue growth and repair (15, 20–26). In addition, we make use of multiple genetic variants in which cells of the immune system are identified with easily visualized markers.

Here, we leverage these strengths to investigate T cell behavior within the zebrafish fin, a highly accessible and regenerating appendage. We have uncovered an unexpected, tissue-specific mode of T cell organization characterized by collective streaming migration that emerges during maturation and precedes tissue vascularization. This behavior is distinct from previously described epidermal T cell surveillance networks and is restricted to T cells, not shared by other immune populations (27). Collective migration is age-dependent and is maintained and re-established following regeneration only in younger, thymus-intact animals, but is delayed and/or lost following thymic involution. Transcriptomic analysis indicates that the composition of both CD4׈ and CD8׈ T cells vary with age and the regenerative state. Furthermore, by employing various genetic models that allow for the modulation of the stromal microenvironment (specifically osteoclasts and macrophages) and by utilizing immunocompromised backgrounds, we have characterized the stromal control over the expression of MHC class I and class II complexes. Together, these findings identify a previously unrecognized, age-regulated mode of tissue-resident adaptive immunity with implications for immune surveillance, regeneration, and age-dependent therapeutic responsiveness.

## Results

### Collective streaming migration of T cells in zebrafish fins

T cells arise in zebrafish during larval stages and persist throughout development (28). We observed a dense population of T cells localized within all five fins of juvenile and adult fish **(Fig. 1A–C, Supplemental Movie 1)**. Previous values have revealed that zebrafish have a total of 2 × 10^5^ total T cells (29). Here, quantification demonstrated a non-negligible number of T cells (approximately 10% of total zebrafish T cells) in the caudal and anal fins (**Fig. 1B**). Live imaging revealed that fin-resident T cells, identified by expression of GFP-*lck*, or mCherry-*lck* undergo coordinated, collective streaming migration that is restricted to the connective tissue of the fins and excluded from the bony rays **(Fig. 1C and Supplemental Movie 2)**. This organized motility was observed across fins, indicating a conserved appendage-specific behavior. Quantitative tracking of individual cells revealed a broad distribution of migration speeds within each fin. Mean velocities ranged from approximately 25 μm/min in the dorsal fin to 40 μm/min in the anal fin (Fig. 1D,E). To characterize migratory behavior, we performed power-law analysis of the mean squared displacement (MSD) (30–32). The slope of the log–log MSD plot (α) distinguishes Brownian (α = 1), subdiffusive (α < 1), and superdiffusive (α > 1) motion (Fig. 1F)(30–32). At the population level, fin-resident T cells exhibited a mixture of subdiffusive and superdiffusive behaviors. Analysis of individual trajectories within the anal fin revealed pronounced heterogeneity within the streaming population **(Fig. 1G–I)**. Representative cells displayed long-range displacements spanning tens of microns over minute timescales, with individual cells alternating between subdiffusive and superdiffusive motility regimes (**Fig. 1I).** Together, these data demonstrate that fin-resident T cells self-organize into dynamic, collectively migrating streams composed of cells with heterogeneous but coordinated motility behaviors.

**Figure 1.**
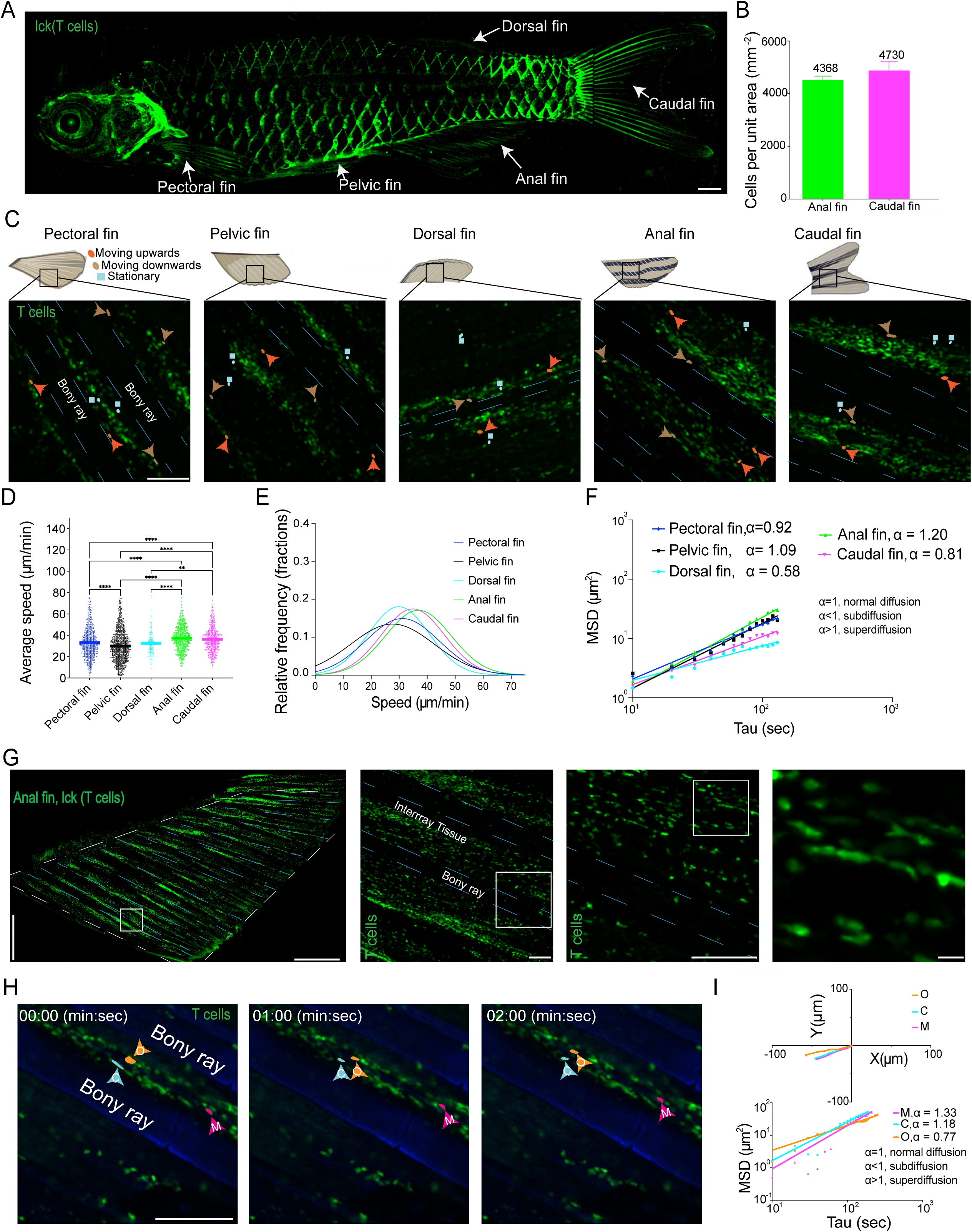
Collective streaming migration of T cells in zebrafish fins. (A) Fluorescence micrograph showing the broad distribution of Tg(*lck:* mCherry, green)+ T cells in an adult fish. Cells are present in the epidermis and densely located within the five fins (arrows), scale bar = 1 mm. (B) Graph comparing the relative surface area covered by T cells per unit area in the anal (average area of anal fin, 13.7 mm^2^) and caudal fins (area of caudal fin, 31.3 mm^2^), N = 3 fish. (C) Micrographs of individual fins displaying the spatial distribution of T cells, which are mostly located within the interray tissue (bony rays represented with the blue broken lines). Highlighted cells indicate differences in motility. Upward-moving cells with orange arrow and shape, downward-moving cells with brown arrow and shape, and searching cells with cyan shape, scale bar = 100 μm. (D) Graph quantifying the average speeds of T cells found in each fin in an individual fish, scale bar = 100 μm. Individual dots correspond to individual cells. (E) Histogram showing that T cell average speeds follow a single Gaussian distribution. (F) Mean square displacement (MSD) curves and fits illustrating differences in T cell motility. (G) Sectioning of the anal fin and the zoom-in images (indicated by the white boxes) of the anal fin, revealing T cells. (H) Micrographs showing individual highlighted cells obtained from time-lapse acquisition. The anal fin of Tg (*lck*:mCherry, green) was stained with calcein to show the bony rays in blue, scale bar = 100 μm. (I) Individual plots of displacement and corresponding MSD curves and fits, used to categorize types of motilities. Statistical analysis was performed using an ordinary one-way ANOVA. *p<0.05, **, p<0.01, *** p<0.001, and **** p<0.0001.

### T cell arrival precedes vascularization of the anal fin

The emergence of coordinated T cell streaming within fin connective tissue prompted us to investigate when and how T cells first colonize the fin, and whether their arrival depends on vascular or lymphatic conduits. The anal fin provides a well-characterized developmental system in which lymphatic vessels represent the first vascular structures and arise at approximately 14 days post-fertilization (dpf) (33). We therefore asked whether T cell infiltration occurs only after lymphatic development. The median fin fold, the precursor to the anal fin, becomes morphologically apparent at 9 dpf, at which stage T cells were not detected. By 11 dpf, Lck T cells were observed arriving and localizing to the median fin fold, well before the onset of lymphatic sprouting. Increasing numbers of Lck T cells populated the nascent anal fin through 13 dpf, prior to vascularization **(Fig. 2A,B).** At these early stages, fin-resident T cells exhibited predominantly random motility indicating that tissue colonization precedes both vascular development and the later emergence of collective migration.

**Figure 2.**
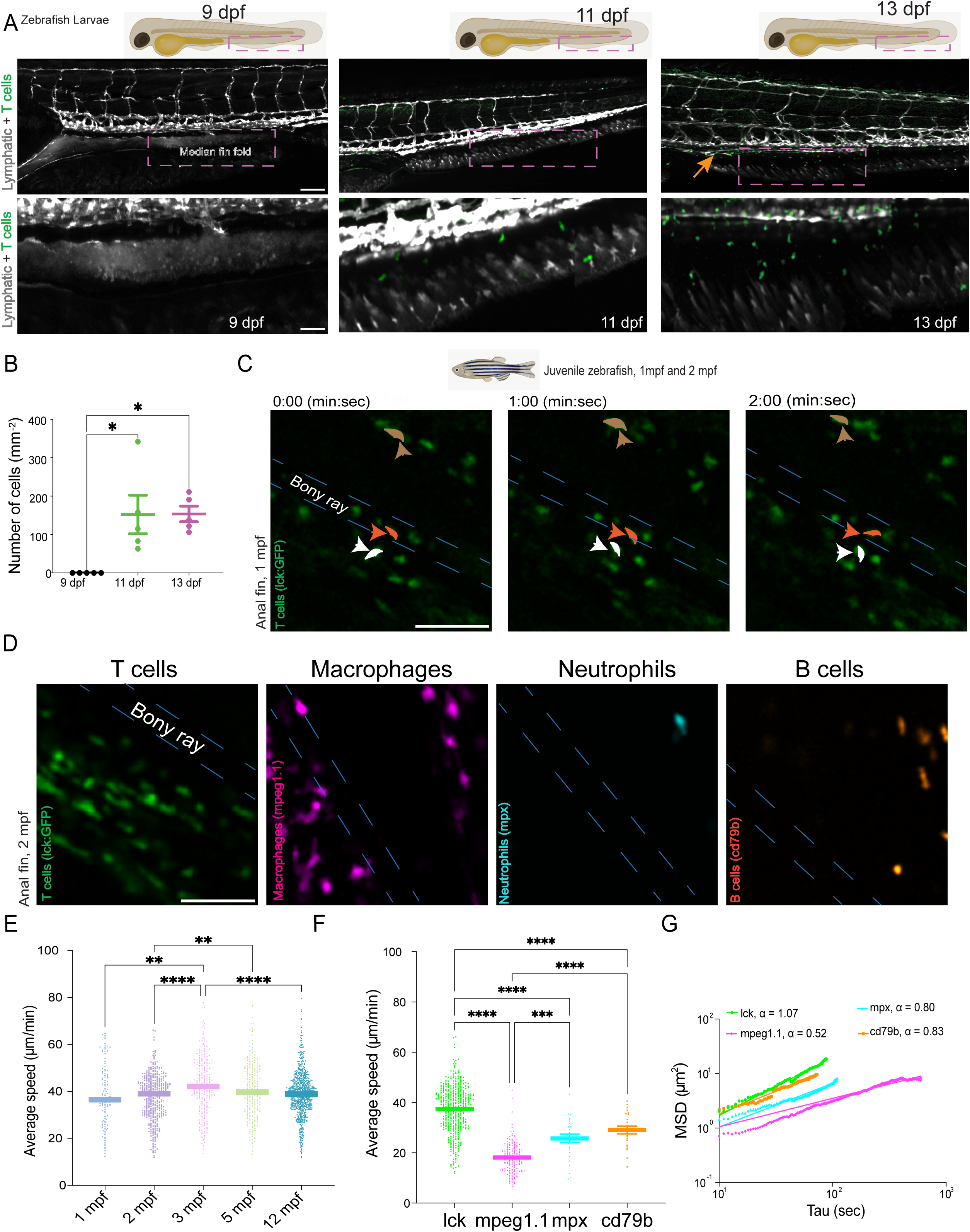
Developmental stage governs the transition from random to collective T cell migration. (A) Schematic and fluorescence micrographs illustrating longitudinal imaging of the median fold (precursor anal fin) in larval zebrafish at 9, 11, and 13 days post fertilization (dpf), showing T cells (*lck*: GFP, green) and lymphatic vessels (*lyve*:dsRed, white), scale bar = 100 μm. (B) Graph showing T cell density (number of T cells per unit area) as a function of development. Each dot represents a single fish. Statistical analysis was performed using an ordinary one-way ANOVA. *p<0.05. (C) Micrographs displaying still images from time-lapse imaging at 1 month post fertilization (mpf), where T cells exhibit random movement within the anal fin and localize in both the bony ray (blue broken lines) and interray tissues (highlighted), scale bar = 50 μm. (D) Micrographs showing the spatial distribution of T cells (*lck*: GFP), Macrophages (*mpeg1.1*:mCherry), Neutrophils (*mpx*: GFP), and B cells (*cd79b*: GFP) in 2 mpf juvenile fish, scale bar = 50 μm. Macrophages are present in both the bony rays and interray tissues, while T cells, Neutrophils, and B cells are restricted to the interray tissue. (E) Quantification of the average speeds of T cells across all ages within the anal fins of 5 fish. Individual dots represent the individual cells. (F) Quantification of the average speeds for all observed immune cell types found in the anal fin, N= 5 fish. (G) Mean Squared Displacement (MSD) curves and fits, used to categorize the motility types for each immune cell population, N = 5 fish. Statistical analysis was performed using an ordinary one-way ANOVA. *p<0.05, **, p<0.01, *** p<0.001, and **** p<0.0001.

### Developmental stage governs the transition from random to collective T cell migration

Anal fin maturation, including the development of skeletal, soft interray, and vascular tissues, was complete by approximately one-month post-fertilization (mpf) (33). At this stage, T cells densely populated the soft interray tissue, with a smaller population also present within the bony rays **(Fig. 2C).** These cells exhibited random motility **(Fig. 2C; Supplemental Movie 3).** A pronounced transition occurred by 2 mpf: T cells were no longer detectable within the bony rays and instead adopted coordinated collective streaming behavior within the soft interray connective tissue **(Fig. 2D; Supplemental Movie 3**). This migratory organization was maintained throughout late juvenile, adult, and old fish. Quantitative analysis revealed that average T cell migration speed peaked at approximately 45 μm/min at 3 mpf and declined with age, reaching average values of 40 μm/min in middle-aged fish at 12 mpf **(Fig. 2E–G).** Analysis of other immune populations within the anal fin identified mpeg׈ macrophages, mpx׈ neutrophils, and B cells. With the exception of macrophages, these cells were largely confined to the soft interray tissue, migrated more slowly, exhibited predominantly sub-diffusive motility, and did not display collective streaming behavior **(Fig. 2D–G; Supplemental Movies 4).**

### Age dependent stromal remodeling accompanies spatial redistribution, and motility shifts of T cells during fin regeneration

These age-dependent changes in immune dynamics prompted us to ask if there are changes in the tissue architecture that change due to aging. Moreover, zebrafish can regenerate multiple organs thus we also could ask how newly infiltrated T cells behave in newly formed tissues. We therefore used the regenerating zebrafish fin to investigate how tissue architecture shapes T cell behavior within newly formed tissue (34). In young adult fish, three months after fertilization, T cells successfully repopulated the regenerated fin following amputation. They also re-established collective streaming migration within tissues that had developed spatial segregation between the fin rays and the interray regions. Quantitative analysis revealed that T cells in regenerated tissue migrated more slowly than those in unamputated fins **(Fig. 3A–D; Supplemental Movie 5),** and a small number of T cells were detected within nascent bony rays. In contrast, fin regeneration in middle aged fish was accompanied by pronounced alterations in T cell spatial organization and motility. Compared to young adults, T cells in older regenerating fins were smaller in size and more broadly dispersed throughout the tissue, rather than being confined primarily to soft interray regions. A substantial fraction of T cells infiltrated newly forming bony rays. Concomitant with this spatial redistribution, T cells exhibited a marked transition from coordinated collective streaming to predominantly random motility **(Fig. 3A–D; Supplemental Movie 5).**

**Figure 3.**
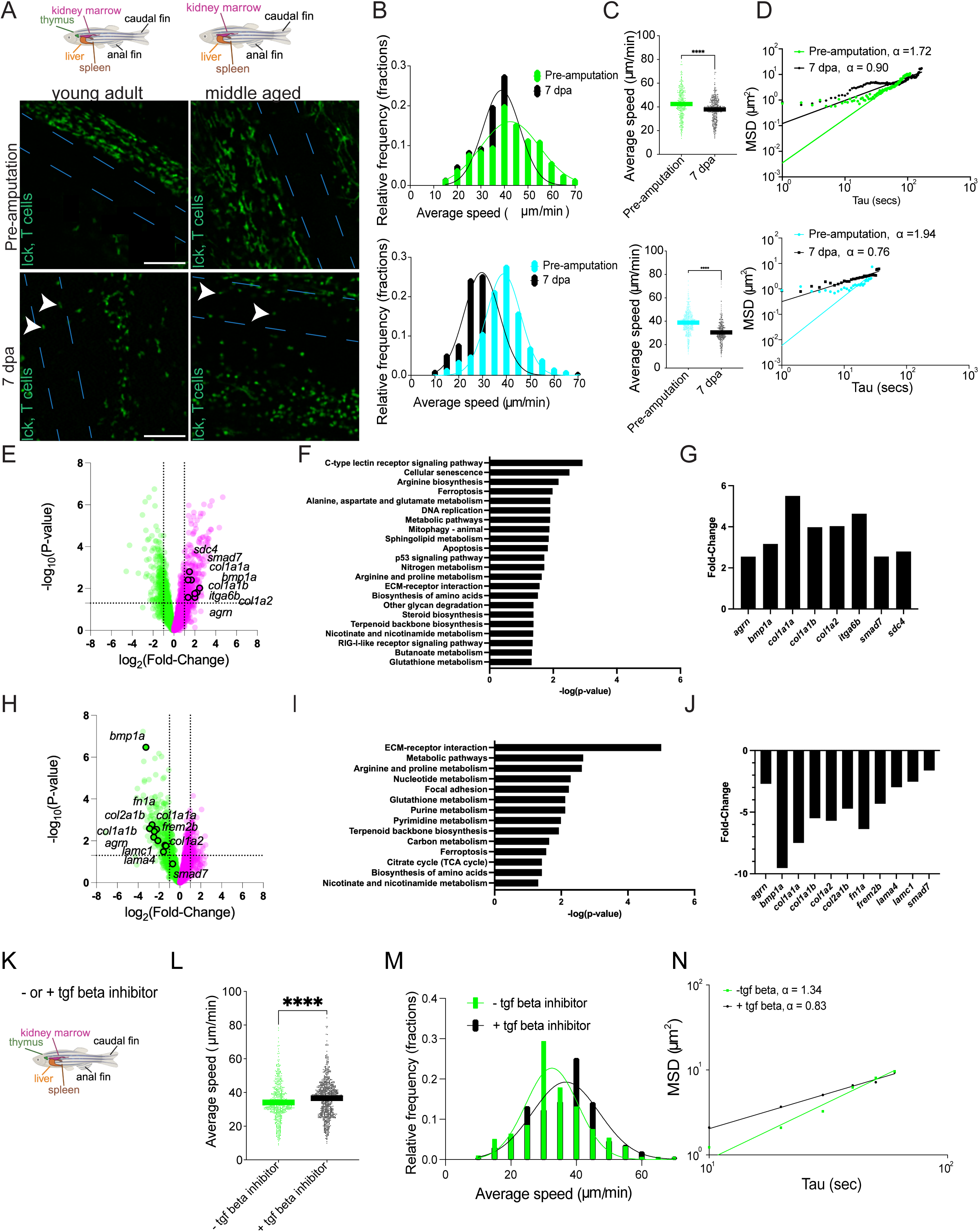
Age dependent stromal remodeling accompanies spatial redistribution, and motility shifts of T cells during fin regeneration. (A) Schematic illustrating thymus involution in young (3 months post-fertilization [mpf]) vs. middle-aged (12 mpf) adult fish. Micrographs show the distribution of *lck*+ T cells in the anal fin pre-and 7 days post-amputation (dpa) for both age groups, scale bar = 50 μm. (B) Histograms showing the distribution of average T cell speeds for young (top) and middle-aged (bottom) fish, N = 5 fish per group. (C) Quantification of average T cell speeds pre-and 7 dpa for young (top) and middle-aged adults (bottom), N = 5 fish per group, Statistical analysis using an unpaired welch t test. *p<0.05, **, p<0.01, *** p<0.001, and **** p<0.0001. (D) Mean Squared Displacement (MSD) curves and fits, used to categorize T cell motility types pre-and 7 dpa for young (top) and middle-aged adults (bottom). (E) Volcano plots displaying differentially expressed genes (DEGs) comparing young (3 mpf) pre-amputation to middle-aged (12 mpf) pre-amputation fish. Magenta dots indicate genes expressed higher in young fish, and green dots indicate genes expressed higher in middle-aged fish. (F) KEGG pathway analysis showing enrichment of pathways related to metabolism, DNA replication, and extracellular matrix (ECM) receptor interaction. (G) Fold changes associated with the ECM receptor pathway comparing young pre-amputation to middle-aged pre-amputation fish. (H) Volcano plots displaying DEGs comparing middle-aged (12 mpf) pre-amputation to middle-aged post-amputation fish. Magenta dots indicate genes expressed higher pre-amputation, and green dots indicate genes expressed higher post-amputation. (I) KEGG pathway analysis showing enrichment of pathways related to metabolism, DNA replication, and ECM receptor interaction comparing middle-aged pre-amputation to middle-aged post-amputation fish. (J) Fold changes associated with the ECM receptor pathway comparing middle-aged pre-amputation to middle-aged post-amputation fish. (K) Schematic of the protocol for treating 3 mpf fish pre-amputation with a TGF-beta inhibitor for 30 minutes. (L) Quantification of the average T cell speeds pre-and post-TGF-beta inhibition for young adults. (M) Histograms showing the distribution of average T cell speeds pre-and post-TGF-beta inhibition for young adults. (N) Mean Squared Displacement (MSD) curves and fits, used to categorize T cell motility types pre-and post-TGF-beta inhibition for young adults. Statistical analysis showing the comparison between the migration speed without (-) and with (+) tgf-beta inhibitor using a two-tailed paired t test. *p<0.05, **, p<0.01, *** p<0.001, and **** p<0.0001.

To identify stromal factors associated with this age dependent regenerative response, we performed comparative transcriptomic analyses of anal fins from young and older fish prior to amputation **(Fig. 3E–G)** and from older fish before and after regeneration **(Fig. 3H–J).** Differential expression analysis revealed distinct gene expression programs across conditions. KEGG pathway analysis identified enrichment of pathways related to metabolism, DNA replication, and extracellular matrix receptor interaction **(Fig. 3F,I).** Notably, collagens and integrins were expressed at higher levels in young fish prior to amputation compared to older fish, whereas these same genes were downregulated following amputation in older fish **(Fig. 3G, J).** We observed that genes associated with bone morphogenesis and TGF-β signaling were upregulated in 3 months post-fertilization (mpf) fish compared to 12 mpf fish prior to amputation (35, 36). Following amputation, these genes were expressed at lower levels in the 12 mpf fins. Given this observation, we investigated whether inhibiting TGF-β would affect cell migration in young adult fish **(Fig. 3K).** Our results indicate that the average speed of cell migration increased following TGF-β inhibition **(Fig. 3L, M, Supplemental Movie 6).** Despite this increase in average speed of migration of T cells, power law analysis of the mean squared displacement (MSD) curves revealed a loss of collective streaming motion with the cells exhibiting sub-diffusive behavior **(Fig. 3N).** Together, these data indicate that age dependent remodeling of the extracellular matrix accompanies impaired restoration of coordinated T cell organization during regeneration with a likely role of TGF-β signaling in this process.

### Age and immune microenvironment regulate antigen presentation programs in regenerating fins

Given the abundance of immune cells within the anal fin, we examined how age and immune microenvironment influence T cell composition and antigen presentation during regeneration **(Fig. 4A).** RNA sequencing was performed on fin tissue and immune relevant organs such as thymus, kidney, spleen, and liver from young adult and middle-aged zebrafish following thymic involution **(Fig. 4A, Supplemental Fig 1-3).** To assess microenvironmental context dependence, we analyzed tissue from zebrafish strains with csf1ra deactivation and animals with reduced T and B cells (SCID) **(Fig. 4B; Supplemental Fig. 1).** Csfr1a is implicated in macrophage and osteoclast regulation in zebrafish whereas SCID fish have reduced numbers of T and B cells (37–41). In immune competent animals, middle aged fish showed elevated expression of T cell associated markers lck, cd4 (helper), and cd8 (cytotoxic) relative to young adults prior to amputation, with persistence of these markers in regenerated fins (**Fig 4B, E-F).** In contrast, cd4 and cd8 expression was significantly reduced in csf1ra deactivated and SCID animals, indicating that fin resident T cell composition is sensitive to immune microenvironmental cues (**Fig 4 E-F).** Lck (T cell) expression was elevated in SCID fish prior to amputation, whereas other immune markers including cd3 (naïve), foxp3a (T-regs), tox, (T cells) and cd79a (T and B cells) did not vary with age or regeneration status **(Fig 4 B, Supplemental Fig. 1).** The T cell proliferation associated marker top2a increased following amputation across all conditions, consistent with regenerative growth **(Fig4 G).** Markers of other immune populations were unchanged **(Figure 4 C-D, Supplemental Fig. 1).** Notably markers of Natural Killer-like cells and Dendritic cells were expressed at extremely low values.

**Figure 4.**
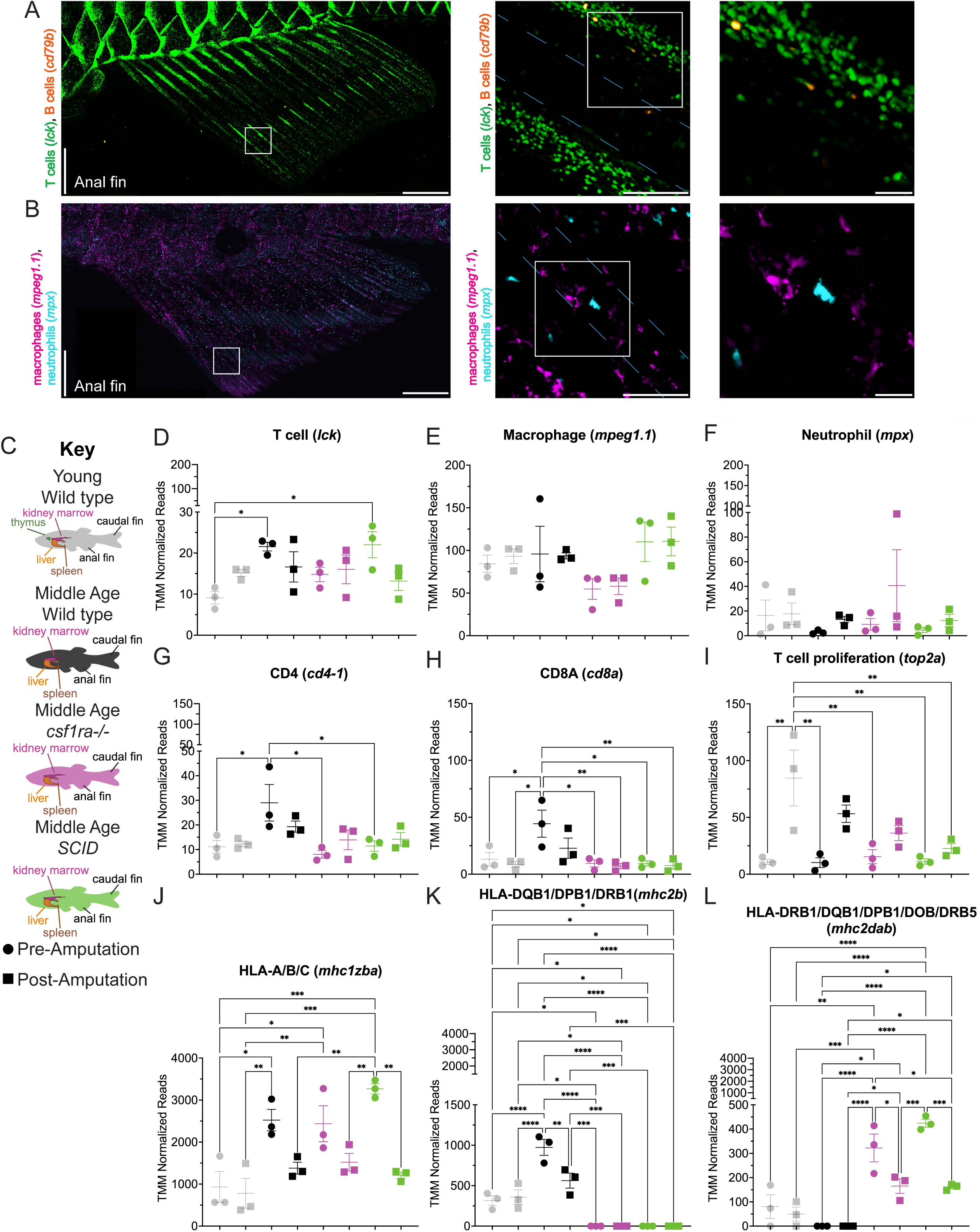
(A) Micrographs show overview of young adult anal fin, where the insets show the relative distributions of *lck*+ T cells (green) and *cd79b* + B cells (orange), scale bar = 1 mm. The white boxes show the zoom-in images of T and B cells remain localized to the interray tissue and absent from the bony rays (white broken lines), scale bar = 100 μm. (B) Micrographs show an overview of the young adult anal fin, where the insets show the relative distributions of *mpeg1.1*+ macrophages (magenta) and *mpx*+ (cyan) neutrophils, scale bar = 1 mm. The white boxes show the zoom-in images of macrophages and neutrophils, scale bar = 100 μm. The white broken lines represent the bony rays. (C) Schematic diagram illustrating the isolation of tissues for transcriptomic analysis, categorized by strain-osteoclasts and macrophage impaired fish (*csf1ra-/-),* reduced T and B cells (SCID), age (before (3mpf) and after thymus involution (12 mpf)), and amputation status (pre and post regeneration). (D-K) Graphs presenting the expression levels of markers for immune cells, T cells, *lck*, (D), Macrophages, *mpeg1.1*, (E), Neutrophils, *mpx*, (F), T cell marker CD4, *cd4-1*, (G), T cell marker CD8, *cd8a*, (H), T cell proliferating marker, *top2a*, (I), and the antigen presentation machinery (J-K). Major histocompatibility complexes: *mhc1zba* (MHC-I ortholog to human HLA-A, HLA-B, HLA-C), *mhc2b* (MHC-II ortholog to human HLA-DQB1, HLA-DPB1, HLA-DRB1) and *mhc2dab* (MHC-II ortholog to human HLA-DRB1, HLA-DQB1, HLA-DPB1, HLA-DOB, HLA-DRB5). Statistical analysis using one-way ANOVA Turkey’s multiple comparisons test where alpha is 0.05 on the mean. *p<0.05, **, p<0.01, *** p<0.001, and **** p<0.0001.

T cells recognize antigens presented by specialized glycoproteins called Major Histocompatibility Complex (MHC) molecules (42–45). These MHC molecules are vital for immune surveillance, enabling the immune system to identify pathogens or abnormal cells and mount an appropriate response to eliminate them (42–45). Analysis of MHC complexes reveals distinct age and context-dependent regulation. Expression of mhc1zba (HLA MHC I) increased with age and decreased following amputation, independent of immune background **(Fig. 4H)**(46). MHC class II components mhc2b and mhc2dab showed greater dependence on the microenvironment (47). Mhc2b expression was undetectable in csf1ra-deactivated and SCID animals, while mhc2dab expression was absent in middle-aged wild-type fins but present in young adults before and after amputation **(Fig 4I-J).** Following regeneration, mhc2dab expression decreased selectively in csf1ra-deactivated and SCID animals, whereas mhc2b expression was reduced in middle-aged fish **(Fig 4I-J).** In parallel, we assessed the systemic changes of immune cell markers in other leucocyte-rich organs, the thymus, kidney, spleen, and liver (**Supplemental Fig. 2-3**). Expression of naïve, CD4, and CD8 T cell markers was highest in young adult fish compared to middle-aged fish. Furthermore, the reduced expression of these T cell markers and mhc2b in csf1ra deactivated and SCID animals provides additional evidence for systemic effects caused by germline mutations. Together, these findings demonstrate that age and immune microenvironment coordinately regulate antigen presentation programs in regenerating tissue.

## Discussion

A fundamental question in immunology is how tissue-resident T cells, a subset of tissue-tropic lymphocytes, are maintained and functionally organized within non-inflamed peripheral tissues across the lifespan (1, 8, 48, 49). In mammals, it remains unclear whether the initial cohort of T cells that seeds peripheral tissues after thymic education is sustained through local self-renewal or replenished by extrathymic sources, particularly following the age-associated involution of the thymus (1, 8, 48). Addressing these longitudinal questions in mammals is technically challenging (50–53). Here, using the zebrafish fin as a tractable in vivo model, we identify a previously unrecognized mode of tissue-specific adaptive immune organization characterized by collective streaming migration of resident T cells and find that this phenomenon is regulated by both the tissue architecture/composition and the maturation of the T cells.

Our findings indicate that tissue-resident T cell organization is developmentally programmed but microenvironmentally instructed. T cells colonize fin tissue early, prior to vascular or lymphatic development, demonstrating that initial tissue residency does not require classical immune conduits (**Figure 2**). However, early-arriving T cells exhibit predominantly random motility and only later transition into coordinated collective streaming as fin architecture matures. This temporal separation suggests that collective organization is not an intrinsic property of T cells alone but rather reflects immune adaptation to a tissue-specific niche. Such niche-dependent organization is consistent with emerging concepts of tissue-resident immunity in which local stromal and architectural cues shape immune behavior and identity (5, 8, 49).

Aging disrupts this organization in two related contexts. First, baseline T cell motility declines with age. Second, following fin amputation, T cells successfully repopulate regenerated fins in both young and middle-aged animals. However, while young adults robustly re-establish collective streaming once tissue architecture is restored, middle-aged, thymic-involuted fish fail to recover coordinated migration. Because T cells repopulate regenerated fins even after thymic involution, these findings indicate that tissue re-entry is not thymus dependent. Recently, it has been shown that zebrafish T regs ( foxp3a+) are instrumental in regeneration of the spinal cord and heart (54). These cells are recruited to sites of injury and tissue specific factors facilitate healing (54). Our results show that there is also comparable expression in foxp3 pre-and post-regenerated fins independently of age and strain. Instead, the failure to re-establish organized streaming in older fish suggests that age-dependent changes in the tissue microenvironment, rather than T cell availability alone, limit restoration of collective organization. Instead, an increased expression of the proliferation-associated marker top2a further supports a model in which regeneration involves both recruitment and local expansion of existing peripheral T cell pools.

Transcriptomic analyses identify stromal remodeling as a prominent feature associated with age-dependent changes in T cell organization. Genes linked to extracellular matrix composition and cell–matrix interactions, including collagens and integrins, display differential regulation with age and following regeneration (55–57). In parallel, components of TGF-β– associated signaling pathways are more highly expressed in young adult fins prior to amputation and reduced in older fins following regeneration. TGF-β signaling is widely recognized as a regulator of extracellular matrix synthesis, deposition, and remodeling during tissue growth and repair, and its activity is influenced by, and can influence, local tissue architecture (35, 36). Consistent with this relationship, pharmacological inhibition of TGF-β in young adult fish altered T cell migratory behavior, increasing average cell speed while coinciding with a loss of coordinated collective streaming. These results also argue for a dual role of the tissue substratum and the T cells themselves in coordinating the streaming motion of the cells, suggesting that the T cells identify an age-dependent component of the tissue architecture along which to move to facilitate the streaming phenomenon.

While these findings do not establish a direct causal role for TGF-β signaling in organizing collective T cell behavior, they suggest that stromal pathways linked to extracellular matrix remodeling may contribute to the tissue contexts that support coordinated T cell organization. Instead, TGF-β also regulates T cell biology. Specifically, it is required for induction of regulatory T cells (Treg) (58). Thus, one idea is that the dysregulation caused by TGF-β inhibition could be because a subset of the resident T cells in the fin is dually affected by inhibition of TGF-β. Further studies will be required to determine how specific stromal signaling pathways interact with tissue architecture to shape immune cell dynamics during regeneration.

In parallel, transcriptomic analyses reveal that aging and immune microenvironment coordinately regulate antigen presentation programs within regenerating tissue. Expression of MHC class I and class II components varied with age, regenerative state, and stromal background, with MHC class II expression showing particular sensitivity to macrophage-and osteoclast-associated cues (**Figure 4**). These observations are consistent with the role of macrophages as professional antigen-presenting cells and with prior work demonstrating that csf1ra macrophages regulate immune cell composition, tissue remodeling, and mechanical properties in zebrafish tissues. Moreover, analysis of the caudal fin following amputation recently revealed conserved cell types (epithelial, hematopoietic, mesenchymal) (59). However, an increase in immunoproteasome subunits was observed specifically in the epithelial and hematopoietic cells (59). The authors speculated that this facilitated acceleration of antigen processing, which could be important for immune cell recruitment.(59). Our data suggests this conservation applies to the anal fin and exhibits age dependence. The involvement of osteoclast-associated pathways further suggests that organ-specific stromal programs contribute to shaping antigen presentation capacity within peripheral tissues. Together, these findings point to a close coupling between stromal composition, immune cell crosstalk, and adaptive immune programming *in situ*. Notably, perturbation of stromal signaling pathways, including TGF-β, disrupts coordinated T cell organization despite preserved or increased motility, indicating that appropriate tissue architecture, rather than cell movement alone, is required for functional tissue-resident immune behaviors.

The collective organization described here is distinct from other known modes of organ-specific T cell surveillance (60, 61). Epidermal tessellated lymphoid networks support body-wide immune coverage at later developmental stages, whereas fin-resident streaming emerges earlier and is restricted to connective tissue compartments (27). Together with prior work demonstrating tissue-specific amoeboid migration strategies, our findings underscore that T cell behavior is highly context dependent, shaped by local developmental history, stromal architecture, and immune composition (17, 27, 60, 61). Within this framework, collective streaming may represent a specialized organizational strategy that supports immune surveillance and coordination within regenerating or mechanically dynamic tissues.

Collectively, these data support a model in which tissue-resident T cells are not passive occupants of peripheral tissues but actively integrate developmental, stromal, and immune cues to establish functional organization across the lifespan. Aging can alter this integration by reshaping antigen presentation landscapes, thereby constraining coordinated immune organization and potentially limiting immune surveillance and regenerative capacity. Here, our findings demonstrate that tissue-specific and systemic microenvironmental cues regulate not only T cell persistence but also T cell organization and functional programming, including antigen presentation capacity, within peripheral tissues. In humans, the age-associated decline in *de novo* T cell production following thymic involution necessitates increasing reliance on the homeostatic maintenance and expansion of the existing naïve T cell pool. Augmenting this diminished reservoir remains a critical therapeutic objective, with current strategies focused on cytokine-and growth factor–mediated support of homeostatic proliferation, hormonal modulation of immune-supportive niches, and adoptive transfer of lymphoid progenitors. In parallel, emerging approaches aimed at recreating immune-supportive environments, including artificial bone marrow or thymus-like grafts, offer promising avenues for restoring adaptive immune competence.

## Materials and methods

All animal experiments were done under protocols, LCB-029 and LCB-031 approved by the National Cancer Institute (NCI) and the National Institutes of Health (NIH) Animal Care and Use Committee.

### Zebrafish husbandry

Zebrafish were maintained at 28.5 °C on a 14-h light/10-h dark cycle according to standard procedures. Larvae were obtained from natural spawning, raised at 28.5 °C, and maintained in fish water, 60 mg sea salt (Instant Ocean, #SS15-10) per liter of DI water. The zebrafish lines used in this study were Tg(*lck*: GFP) in a TAB5 background, the double transgenic Tg (*mpeg1.1*:mCherry; *mpx*:GFP), the Tg(*lyve1*:dsRed) in a EK background, the panther^j4e2^ line (csf1ranull), the prkdcD3612fs line (SCID fish), the Tg(*cd79b*:GFP; *lck*:mCherry) line, and the Tg(*cd79a*:GFP) line. A double transgenic line with T cells and lymphatics was generated by crossing Tg(*lck*:GFP) with Tg(lyve1:dsRed). The Tg(*lck*:GFP) line was a generous gift from the Sood Lab. The Tg(lyve1:dsRed) was a generous gift from Brent Weinstein. The panther^j4e2^ line was a generous gift from David Parichy. The prkdc^D3612fs^ line was a generous gift from David Langenau. The Tg(*cd79b:*GFP;*lck*:mCherry) and Tg(*cd79a*:GFP) were a generous gift from J. Kimble Frazer. Embryos were maintained in E3 media (5 mM NaCl, 0.44 mM CaCl_2_, 0.33 mM MgSO_4_, 0.17 mM KCl, 0.025 mM NaOH, 0.0005% methylene blue), at 28.5°C in tissue culture-treated Petri dishes in a laboratory incubator. Fish were introduced to the fish system at 5 days post fertilization.

### Intravital imaging of zebrafish

Zebrafish were anesthetized using 0.4% buffered tricaine methanesulfonate (Syndel SYNCAINE Cat. No. TRS1). Zebrafish larvae were anesthetized and immobilized in a lateral orientation in 1% (w/v) low gelling temperature agarose (Cat. No. A9414-25G, Millipore Sigma) dissolved in 1X E3 media or laterally oriented in a 60 mm glass-bottom dish (Martek Corporation P60G-0-45-F). To enable high-resolution confocal imaging of mounted larvae, fish were laterally oriented in coverglass-bottom chamber slides (Nunc Lab-Tek Chambered #1.0 Borosilicate Coverglass slides, Cat. No 155383, Thermo Fisher Scientific). 1X E3 media supplemented with 0.4% buffered tricaine was then added to the chamber slides to keep the larvae anesthetized over the course of the experiment. To acquire images of immune cells at single-cell resolution, chamber slides containing the larvae were imaged on Zeiss 780 laser scanning or Nikon A1R confocal microscopes. Three-dimensional tile scans of the entire fish, the anal fin, and the caudal fins were obtained at juvenile and young adult stages. Images were taken at axial steps of 5 μm and stacked to create three-dimensional images. For the time-lapse imaging, One-photon, confocal 12-bit images were acquired with a Zeiss 20x EC Plan-Apochromat, 0.8 NA objective and a digital zoom of 1 or 1.5, resulting in a field of view of 425.1 μm x 425.1 μm or 282.8 μm x 282.8 μm, respectively, for each tile of the image. Pinhole diameter was set at 90 μm for Tg(*lck*:GFP) fish and 149.5 μm for Tg (*cd79b*: GFP; *lck*:mCherry). Samples were simultaneously excited with 488 nm light from an argon laser with a total power of 25 mW, and 561 nm light from a solid-state laser with a total power of 20 mW. Transmitted light was also recorded. Images were taken on two tracks to minimize signal overlap. All lasers were set at or below 18 % total power. A beam splitter, MBS 488/561, was employed in the emission pathway to delineate the red (band-pass filters ∼580-645 nm), and green (band-pass filters ∼493-574 nm) channels. The master gain was set at or below 700 for each channel. The zebrafish larvae were maintained at 28°C for the course of imaging. Pixel dwell times of 1.58 μs were used. Before imaging, adult zebrafish were anesthetized with 0.08 mg/mL tricaine (MS-222/ethyl 3-aminobenzoate) and placed in a 60 mm glass-bottom dish (Martek Corporation P60G-0-45-F), and the anal fin was extended in the dish with a superfine eyelash (Ted Pella, 113) to allow proper imaging. Imaging was performed at 1 second interval for 10 minutes. The overviews of adult whole fish and anal fin were taken with Nikon A1R confocal microscopy with z stack of 5 μm, using a Nikon Plan Apo Lambada 2x for the whole fish and 10x for the anal fin and 20x for the zoom-in images, 0.1 NA objective and a digital zoom of 1 for the whole fish and anal fin or 3 for the zoom-in images, with 1024 x 1024 pixels (1768 μm x 1768 μm) for each tile of 12-bit images. The pinhole diameter was set to 2 μm. The fish was simultaneously excited with 488-nm and 561 nm light from a solid-state laser with a total power of up to 70 mW and 568-nm light from a diode-pumped solid-state laser with a total power of up to 70 mW. All lasers were set at or below 5% total power with the master gain at or below 30. After imaging, the fish were either sacrificed or recovered in fresh system water.

### Intravital immune cell tracking

Time-lapse microscopy images were exported to Imaris File Converter (Imaris File Converter x64 10.2.0), and the converted images were analyzed with the Imaris software (Oxford Instruments, v.10.2.0). Background subtraction was performed prior to cell detection. The cells were detected and tracked using the Cells tracking module in Imaris. The estimated sphere diameter was set at 6 µm for the *lck* and *cd79b* cells, 8 µm for the *mpeg1.1* cells, and 7 µm for the *mpx* cells, and background subtraction was enabled. Tracking was performed using the autoregressive motion model, with a maximum distance of 3 µm and a maximum gap size of 3 frames. The tracks shorter than 30 seconds were excluded from the analysis. Following automated tracking, ten cells were randomly selected and manually inspected for tracking accuracy. Only verified tracks were included in mean squared displacement (MSD) analysis. Track parameters, including mean cell speed (in µm/sec) and cell two-dimensional coordinates (X and Y) at each time point, were exported from Imaris and processed in Microsoft Excel. The track speed was converted to µm/min by multiplying the speed in µm/sec by 60. MSD was calculated from two-dimensional coordinate data using the following equation:

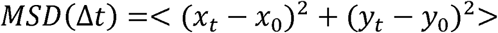

Where x_t_ and y_t_ represent the cell position at time t, x_0_ and y_0_ represent the initial position, and the angle brackets indicate averaging across time points. For each cell, squared displacements in the X and Y directions relative to the initial position were summed to obtain MSD values, which were plotted as a function of time using GraphPad Prism (v.10.0.0). MSD analysis was used as a quantitative measurement of cell motility, where higher MSD values over time correlated to increased migratory capacity and reduced spatial confinement.

## Fish amputation

Anal fins were amputated from fish at 3 and 12 months post-fertilization using a scalpel (Techno cut scalpel #11) while the fish were under anesthesia. Before and seven days after amputation, the fins were imaged as described previously. Following the final imaging, the fish were euthanized. Fish were returned to the fish system after the amputation procedure.

For the TGF-β experiment, the TGF-β inhibitor, LY2157299, generously donated by Dr. Wakefield’s Lab, was diluted to 100 nM in 1X E3 media from a 10 mM stock. The 100 nM dilution was added to the extended anal fin after image acquisition. Imaging was performed after 30 minutes.

### RNA-Sequencing

To assess transcriptomics, individual organs were dissected from freshly euthanized fish and harvested directly into Trizol by Invitrogen (ThermoFisher Scientific # 15596026) and maintained in -20°C for short-term storage until further processing was performed. RNA was extracted using the manufacturer’s protocol. Total RNA was submitted for RNA-Sequencing to the NCI CCR Genomics Core. The libraries were made using Poly(A) Selection Kit (New England Biolabs #E7490L), Dual Index Barcodes (New England Biolabs #E6440S), and Directional RNA Library Kit (New England Biolabs #E7760L). 150bp fragments were sequenced by paired-end 75bp utilizing Illumina NextSeq 2000 P4 XLEAP-SBS Reagent Kit, 200 Cycles (Illumina #20100993). Reads were aligned to GRCz11/danRer11 using HISAT 2. Reads were quantified to annotation model using PartekFlow. Gene counts were normalized using TMM and comparisons between samples were made using ANOVA. KEGG Pathway analysis was done in PartekFlow after filtering transcripts based on ANOVA analysis of Fold Change greater than 2 or less than -2 and a p-value of less than 0.05.

## Acknowledgements

This research was supported by the Intramural Research Program of the National Institutes of Health (NIH). The contributions of the NIH authors were made as part of their official duties as NIH federal employees, are in compliance with agency policy requirements, and are considered Works of the United States Government. However, the findings and conclusions presented in this paper are those of the authors and do not necessarily reflect the views of the NIH or the U.S. Department of Health and Human Services.

This work utilized the computational resources of the NIH HPC Biowulf cluster. (http://hpc.nih.gov) Services were provided by the CCR Genomics Core at National Cancer Institute. All schematics are created with <u>BioRender.com</u>.

## References

1. B. V. Kumar, T. J. Connors, D. L. Farber, Human T Cell Development, Localization, and Function throughout Life. Immunity 48, 202–213 (2018).

2. R. Sender et al., The total mass, number, and distribution of immune cells in the human body. Proc Natl Acad Sci U S A 120, e2308511120 (2023).

3. C. Dominguez Conde et al., Cross-tissue immune cell analysis reveals tissue-specific features in humans. Science 376, eabl5197 (2022).

4. A. K. Simon, G. A. Hollander, A. McMichael, Evolution of the immune system in humans from infancy to old age. Proc Biol Sci 282, 20143085 (2015).

5. P. A. Szabo et al., Single-cell transcriptomics of human T cells reveals tissue and activation signatures in health and disease. Nat Commun 10, 4706 (2019).

6. P. Brodin, M. M. Davis, Human immune system variation. Nat Rev Immunol 17, 21–29 (2017).

7. Y. Wang, C. Dong, Y. Han, Z. Gu, C. Sun, Immunosenescence, aging and successful aging. Front Immunol 13, 942796 (2022).

8. J. J. Thome et al., Spatial map of human T cell compartmentalization and maintenance over decades of life. Cell 159, 814–828 (2014).

9. E. C. Butcher, L. J. Picker, Lymphocyte homing and homeostasis. Science 272, 60–66 (1996).

10. J. E. Park et al., A cell atlas of human thymic development defines T cell repertoire formation. Science 367 (2020).

11. R. E. Niec, A. Y. Rudensky, E. Fuchs, Inflammatory adaptation in barrier tissues. Cell 184, 3361–3375 (2021).

12. S. J. Carmona et al., Single-cell transcriptome analysis of fish immune cells provides insight into the evolution of vertebrate immune cell types. Genome Res 27, 451–461 (2017).

13. E. R. Jerison, S. R. Quake, Heterogeneous T cell motility behaviors emerge from a coupling between speed and turning in vivo. Elife 9 (2020).

14. K. Kissa et al., Live imaging of emerging hematopoietic stem cells and early thymus colonization. Blood 111, 1147–1156 (2008).

15. D. M. Langenau et al., In vivo tracking of T cell development, ablation, and engraftment in transgenic zebrafish. Proc Natl Acad Sci U S A 101, 7369–7374 (2004).

16. S. A. Renshaw, N. S. Trede, A model 450 million years in the making: zebrafish and vertebrate immunity. Dis Model Mech 5, 38–47 (2012).

17. T. F. Robertson et al., Live imaging in zebrafish reveals tissue-specific strategies for amoeboid migration. Development 152 (2025).

18. A. Hasan et al., Dynamic Changes in Lymphocyte Populations Establish Zebrafish as a Thymic Involution Model. J Immunol 212, 1733–1743 (2024).

19. G. Park, C. A. Foster, M. Malone-Perez, A. Hasan, J. J. Macias, J. K. Frazer, Diverse Epithelial Lymphocytes in Zebrafish Revealed Using a Novel Scale Biopsy Method. J Immunol 213, 1902–1914 (2024).

20. M. Gemberling, T. J. Bailey, D. R. Hyde, K. D. Poss, The zebrafish as a model for complex tissue regeneration. Trends Genet 29, 611–620 (2013).

21. S. P. Hui, K. Sugimoto, D. Z. Sheng, K. Kikuchi, Regulatory T cells regulate blastemal proliferation during zebrafish caudal fin regeneration. Front Immunol 13, 981000 (2022).

22. C. D. Paul et al., Tissue Architectural Cues Drive Organ Targeting of Tumor Cells in Zebrafish. Cell Syst 9, 187–206 e116 (2019).

23. C. D. Paul et al., Human macrophages survive and adopt activated genotypes in living zebrafish. Sci Rep 9, 1759 (2019).

24. F. Ellett, L. Pase, J. W. Hayman, A. Andrianopoulos, G. J. Lieschke, mpeg1 promoter transgenes direct macrophage-lineage expression in zebrafish. Blood 117, e49–56 (2011).

25. J. R. Mathias, B. J. Perrin, T. X. Liu, J. Kanki, A. T. Look, A. Huttenlocher, Resolution of inflammation by retrograde chemotaxis of neutrophils in transgenic zebrafish. J Leukoc Biol 80, 1281–1288 (2006).

26. S. A. Renshaw, C. A. Loynes, D. M. Trushell, S. Elworthy, P. W. Ingham, M. K. Whyte, A transgenic zebrafish model of neutrophilic inflammation. Blood 108, 3976–3978 (2006).

27. T. F. Robertson et al., A tessellated lymphoid network provides whole-body T cell surveillance in zebrafish. Proc Natl Acad Sci U S A 120, e2301137120 (2023).

28. D. M. Langenau, L. I. Zon, The zebrafish: a new model of T-cell and thymic development. Nature Reviews Immunology 5, 307–317 (2005).

29. R. Covacu et al., System-wide Analysis of the T Cell Response. Cell Rep 14, 2733–2744 (2016).

30. T. H. Harris et al., Generalized Lévy walks and the role of chemokines in migration of effector CD8+ T cells. Nature 486, 545–548 (2012).

31. R. Metzler, J.-H. Jeon, A. G. Cherstvy, E. Barkai, Anomalous diffusion models and their properties: non-stationarity, non-ergodicity, and ageing at the centenary of single particle tracking. Physical Chemistry Chemical Physics 16, 24128–24164 (2014).

32. R. Metzler, J. Klafter, The random walk’s guide to anomalous diffusion: a fractional dynamics approach. Physics Reports 339, 1–77 (2000).

33. R. N. Das et al., Generation of specialized blood vessels via lymphatic transdifferentiation. Nature 606, 570–575 (2022).

34. M. Westerfield, The zebrafish book : a guide for the laboratory use of zebrafish (Brachydanio rerio) (M. Westerfield, Eugene, OR, ed. 2.1., 1994).

35. J. Massague, TGFbeta signalling in context. Nat Rev Mol Cell Biol 13, 616–630 (2012).

36. L. Richardson, S. G. Wilcockson, L. Guglielmi, C. S. Hill, Context-dependent TGFbeta family signalling in cell fate regulation. Nat Rev Mol Cell Biol 24, 876–894 (2023).

37. M. Chatani, Y. Takano, A. Kudo, Osteoclasts in bone modeling, as revealed by in vivo imaging, are essential for organogenesis in fish. Dev Biol 360, 96–109 (2011).

38. I. H. Jung et al., Impaired Lymphocytes Development and Xenotransplantation of Gastrointestinal Tumor Cells in Prkdc-Null SCID Zebrafish Model. Neoplasia 18, 468–479 (2016).

39. D. M. Parichy, D. G. Ransom, B. Paw, L. I. Zon, S. L. Johnson, An orthologue of the kit-related gene fms is required for development of neural crest-derived xanthophores and a subpopulation of adult melanocytes in the zebrafish, Danio rerio. Development 127, 3031–3044 (2000).

40. P. Herbomel, B. Thisse, C. Thisse, Zebrafish early macrophages colonize cephalic mesenchyme and developing brain, retina, and epidermis through a M-CSF receptor-dependent invasive process. Dev Biol 238, 274–288 (2001).

41. W. Y. So et al., Macrophages Mediate Mesoscale Brain Mechanical Homeostasis. Adv Mater 10.1002/adma.202517493, e17493 (2025).

42. N. Pishesha, T. J. Harmand, H. L. Ploegh, A guide to antigen processing and presentation. Nat Rev Immunol 22, 751–764 (2022).

43. J. Neefjes, M. L. Jongsma, P. Paul, O. Bakke, Towards a systems understanding of MHC class I and MHC class II antigen presentation. Nat Rev Immunol 11, 823–836 (2011).

44. A. D. Waldman, J. M. Fritz, M. J. Lenardo, A guide to cancer immunotherapy: from T cell basic science to clinical practice. Nat Rev Immunol 20, 651–668 (2020).

45. M. Wieczorek et al., Major Histocompatibility Complex (MHC) Class I and MHC Class II Proteins: Conformational Plasticity in Antigen Presentation. Front Immunol 8, 292 (2017).

46. H. Dirscherl, S. C. McConnell, J. A. Yoder, J. L. de Jong, The MHC class I genes of zebrafish. Dev Comp Immunol 46, 11–23 (2014).

47. H. Ono et al., Major histocompatibility complex class II genes of zebrafish. Proc Natl Acad Sci U S A 89, 11886–11890 (1992).

48. N. Lam et al., Asynchronous aging and turnover of human circulating and tissue-resident memory T cells across sites. Immunity 58, 2271–2288 e2276 (2025).

49. K. Okla, D. L. Farber, W. Zou, Tissue-resident memory T cells in tumor immunity and immunotherapy. J Exp Med 218 (2021).

50. N. Lam, Y. Lee, D. L. Farber, A guide to adaptive immune memory. Nat Rev Immunol 24, 810–829 (2024).

51. D. Schienstock, S. N. Mueller, Moving beyond velocity: Opportunities and challenges to quantify immune cell behavior. Immunological Reviews 306, 123–136 (2022).

52. R. N. Germain, E. A. Robey, M. D. Cahalan, A decade of imaging cellular motility and interaction dynamics in the immune system. Science 336, 1676–1681 (2012).

53. M. Alieva, L. Ritsma, R. J. Giedt, R. Weissleder, J. van Rheenen, Imaging windows for long-term intravital imaging: General overview and technical insights. Intravital 3, e29917 (2014).

54. S. P. Hui et al., Zebrafish Regulatory T Cells Mediate Organ-Specific Regenerative Programs. Dev Cell 43, 659–672 e655 (2017).

55. I. J. Marques, E. Lupi, N. Mercader, Model systems for regeneration: zebrafish. Development 146 (2019).

56. K. S. Midwood, L. V. Williams, J. E. Schwarzbauer, Tissue repair and the dynamics of the extracellular matrix. Int J Biochem Cell Biol 36, 1031–1037 (2004).

57. J. Shen et al., Live visualization of extracellular matrix dynamics during development and regeneration in zebrafish. bioRxiv 10.1101/2025.11.02.686082, 2025.2011.2002.686082 (2025).

58. J. M. Moreau, M. Velegraki, C. Bolyard, M. D. Rosenblum, Z. Li, Transforming growth factor-beta1 in regulatory T cell biology. Sci Immunol 7, eabi4613 (2022).

59. Y. Hou et al., Cellular diversity of the regenerating caudal fin. Sci Adv 6, eaba2084 (2020).

60. A. Chauveau et al., Visualization of T Cell Migration in the Spleen Reveals a Network of Perivascular Pathways that Guide Entry into T Zones. Immunity 52, 794–807.e797 (2020).

61. J. L. Hor, R. N. Germain, Intravital and high-content multiplex imaging of the immune system. Trends in Cell Biology 32, 406–420 (2022).

