## Supplemental Figure Legend for "Age dependent collective trafficking of tissue resident T cells in zebrafish"

**Supplemental Figure 1**. **Transcriptomic analysis the anal fins of wild-type fish (young and middle-aged), csf1ra-/- and SCID fish pre- and post-amputation.**

(A)Schematic diagram illustrating the isolation of tissues for transcriptomic analysis, categorized by strain- osteoclasts and macrophage impaired fish (*csf1ra-/-*), reduced T and B cells (*SCID*), age (before (3mpf, young wild type) and after thymus involution (12 mpf, middle age wild type) and amputation status (pre- and post-amputation). (B-K) Graphs presenting the expression levels of markers for immune cells, T cells, *cd3eap*, (B), T cell exhaustion marker, *tox* (C), B-cell marker, *cd79a* (D), regulatory T cell marker, *foxp3a*, (E), T and B cell development markers, *rag 1* (F), and *rag 2* (G), natural killer cell marker, *nk1.1*, (H), myeloid and lymphoid cell development marker, *irf8*, (I), early myeloid precursor cell marker, *bm2*, (J), and macrophage marker, *csf1ra* (K). Statistical analysis using one-way ANOVA, Turkey's multiple comparisons test, where alpha is 0.05 on the mean. *p<0.05, **, p<0.01, *** p<0.001, and **** p<0.0001.

**Supplemental Figure 2. Quantification of T cell-specific marker transcriptome across multiple organs in wild-type (young and middle-aged), *csf1ra-/-*, and *SCID* fish in control and post-amputation.** (A) Schematic diagram illustrating the isolation of tissues for transcriptomic analysis, categorized by strain- osteoclasts and macrophage impaired fish (*csf1ra-/-*), reduced T and B cells (*SCID*), age (before (3mpf, young wild type) and after thymus involution (12 mpf, middle age wild type)), and amputation status (pre and post regeneration).

(B-K) Graphs presenting the expression levels of markers for immune cells, T cells, *cd3eap*, (B), and cytotoxic T cell marker, *cd8a*, (C). Statistical analysis using one-way ANOVA, Turkey's multiple comparisons test, where alpha is 0.05 on the mean. *p<0.05, **, p<0.01, *** p<0.001, and **** p<0.0001. N = 3 fish, control fish was not amputated.

**Supplemental Figure 3. Quantification of T cell, antigen presentation, and macrophage-specific markers across multiple organs in wild-type (young and middle-aged), *csf1ra-/-*, and *SCID* fish in control and post-amputation.** (A) Schematic diagram illustrating the isolation of tissues for transcriptomic analysis, categorized by strain- osteoclasts and macrophage impaired fish (*csf1ra-/-*), reduced T and B cells (*SCID*), age (before (3mpf, young wild type) and after thymus involution (12 mpf, middle age wild type)), and amputation status (pre and post regeneration). (B-K) Graphs presenting the expression levels of markers for immune cells, T cells, *lck*, (B), and T helper cell marker, *cd4-1*, (C), antigen-presentation marker expressed by profession antigen presenting cells, *mhc2a*, (D), and macrophage marker, *csf1ra*, (E). Statistical analysis using one-way ANOVA, Turkey's multiple comparisons test, where alpha is 0.05 on the mean. *p<0.05, **, p<0.01, *** p<0.001, and **** p<0.0001.
