## Supplementary figures and images for "Age dependent collective trafficking of tissue resident T cells in zebrafish"

### Supplemental Figures

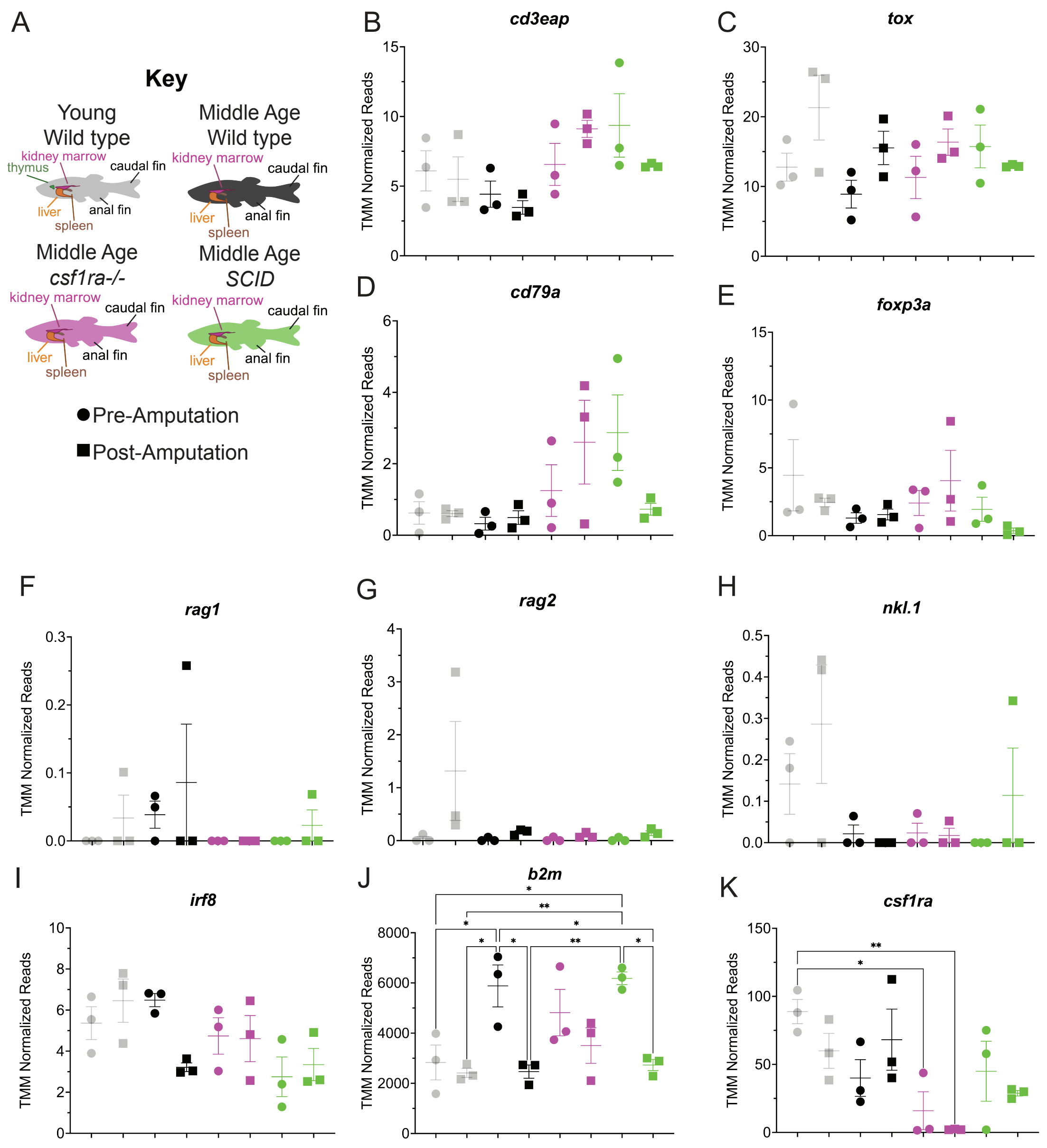


**Supplemental Figure 1**

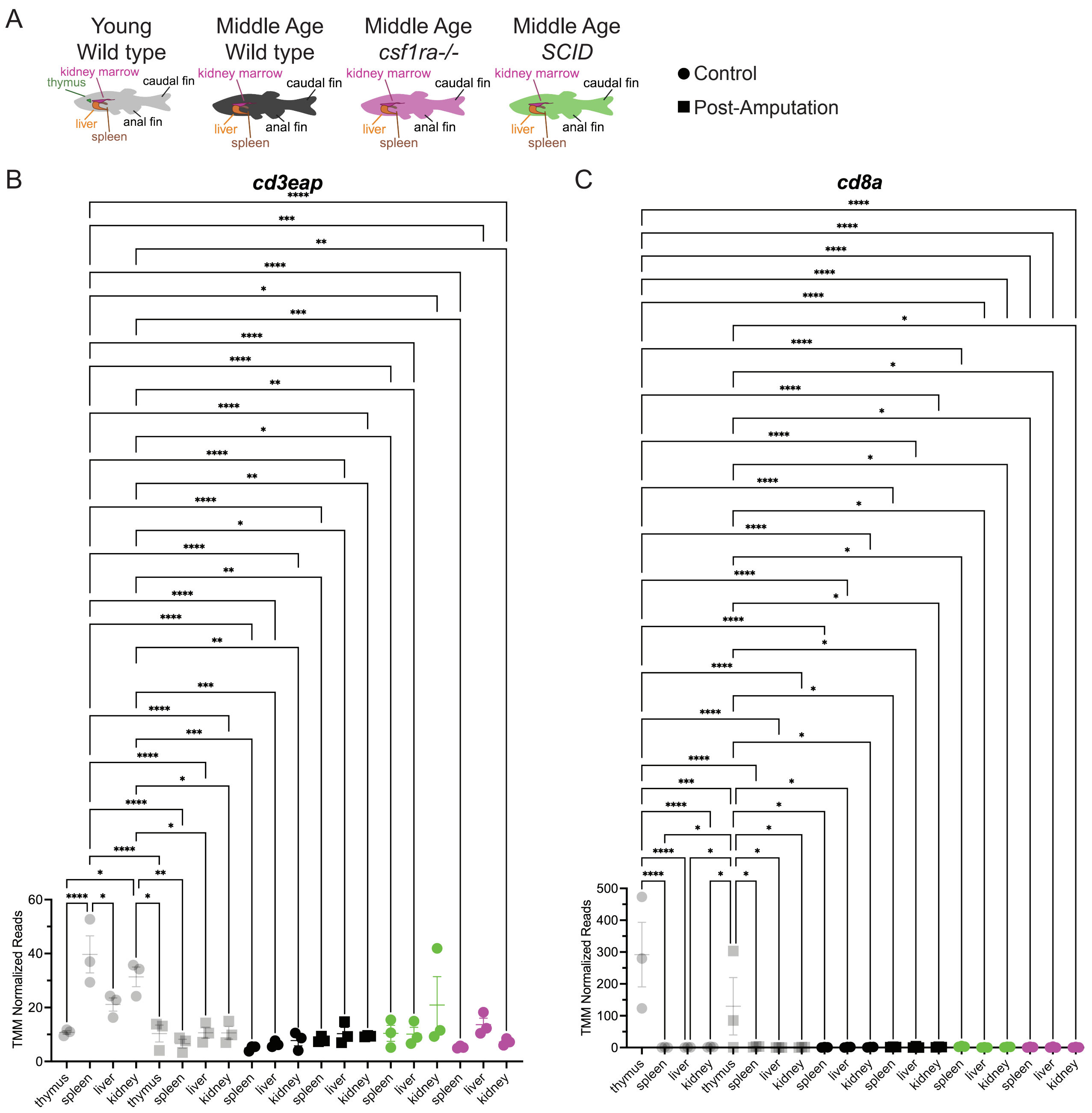


**Supplemental Figure 2**

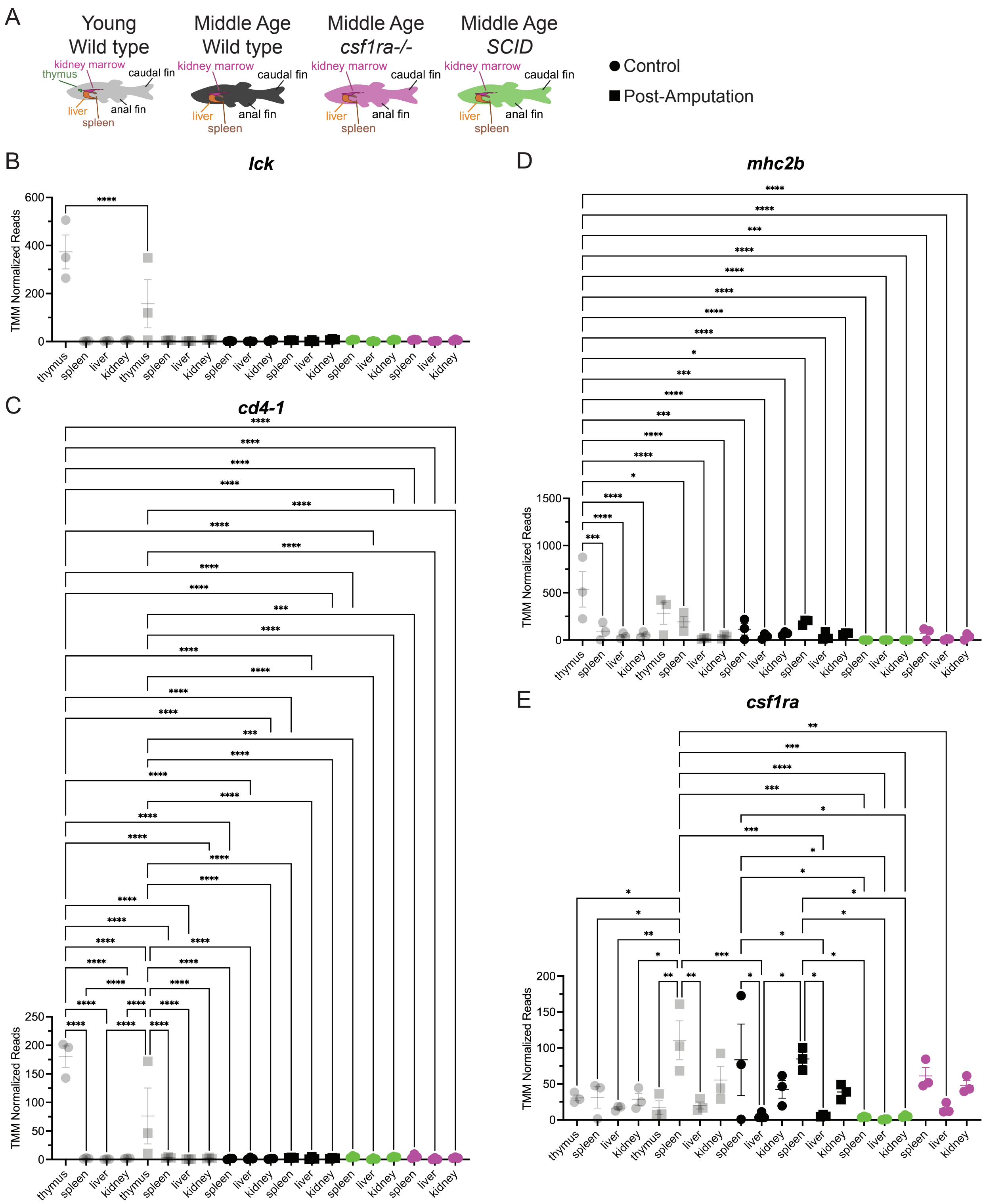


**Supplemental Figure 3**
