## Supplemental Movie Legend for "Age dependent collective trafficking of tissue resident T cells in zebrafish"

**Supplemental movie 1**: Collective migration of T cells in the pectoral, pelvic, dorsal, anal, and caudal fins of a juvenile zebrafish Tg(*lck*:mCherry) shown in green. Live confocal imaging was performed with 20× objective, every 1 second for 10 minutes. The videos are from a single Z plane. These movies correspond to the micrographs in Figure 1C. The frame rate is 320 fps.

**Supplemental movie 2**: Collective migration of T cells in the interray tissue of the anal fin in a juvenile zebrafish Tg(*lck*: mCherry). T cells are shown in green, and the bony rays are shown in blue. Calcein staining was performed to detect the bony rays. Live confocal imaging was performed with 20× objective, every 1 second for 10 minutes. The videos are from a single Z plane. These movies correspond to the micrographs in Figure 1H. The frame rate is 320 fps.

**Supplemental movie 3**: Collective migration of T cells in the interray tissue of the anal fin in zebrafish of different ages, 1-month post-fertilization (mpf), 2 mpf, 3 mpf, 5 mpf, and 12 mpf. Live confocal imaging was performed with 20× objective, every 1 second for 10 minutes. The videos are from a single Z plane. These movies correspond to the graph in Figure 2E. The frame rate is 320 fps.

**Supplemental movie 4**: Differential migration of T cells (green), macrophages (magenta), neutrophils (cyan), and B-cells (orange) in the anal fin of 2 mpf zebrafish. Live confocal imaging was performed with 20× objective, every 1 second for 10 minutes. The videos are from a single Z plane. These movies correspond to the micrographs in Figure 2D. The frame rate is 320 fps.

**Supplemental movie 5**: Collective migration of T cells in the anal fin of young (3 mpf, top panel) and middle-aged fish (12 mpf, bottom panel) zebrafish before (left panel) and after (right panel) amputation. Images were acquired with 20× objective, every 1 second for 10 minutes. The videos are from a single Z plane. These movies correspond to the micrographs in Figure 3A. The frame rate is 320 fps.

**Supplemental movie 6**: Collective migration of T cells in the anal fin of young (3 mpf) zebrafish before (left panel) and after (right panel) adding 100 nM of the TGF-β inhibitor, LY2157299. Live confocal imaging was performed with 20× objective, every 1 second for 10 minutes. The videos are from a single Z plane. These movies correspond to the schematics in Figure 3K. The frame rate is 320 fps.
